# TickMapKB: A FAIR Spatial Knowledgebase of Tick Species and Associated Pathogens in India

**DOI:** 10.64898/2026.06.22.733803

**Authors:** Shreyes Rajan Madgaonkar, Shrish Vashishth, Elango Ayyanar, Srikanth Srirama, Areejit Samal

**Author notes:** Corresponding author (A. Samal). Address for correspondence: Areejit Samal Computational Biology Group, The Institute of Mathematical Sciences (IMSc), CIT Campus, Taramani, Chennai 600113 India.

## Abstract

Ticks transmit pathogens across wildlife, domestic animals, and humans, and are therefore considered important vectors under the One Health framework. In India, the burden of tick-borne diseases, such as Kyasanur Forest Disease and Crimean-Congo Hemorrhagic Fever, remains poorly quantified due to fragmented surveillance and limited spatial data. Here, we present TickMapKB, a curated spatial knowledgebase documenting 72 tick species across more than 600 georeferenced locations in India. The majority of the species belonged to the genera *Haemaphysalis*, *Rhipicephalus*, and *Hyalomma*. Additionally, the resource integrates host associations, pathogen and disease information, morphological identification keys, acaricide resistance profiles, and protein annotations into a single, interactive platform accessible at https://cb.imsc.res.in/tickmapkb/. Specifically, the resource captures 53 pathogens, over 3000 protein annotations, and morphological keys from 26 published resources. TickMapKB thus provides researchers, clinicians, and other stakeholders with integrated spatial, biological, and resistance information to support surveillance planning, resistance management, and tick-borne disease risk assessment.

## Background and Summary

Ticks are obligate parasites that feed on the blood of multiple hosts, including humans^1,2^. Due to their broad host range, prolonged feeding behavior, and ecological adaptability, ticks are among the most important vectors of human and animal pathogens^3,4^. Further, changes in anthropogenic land use and climate-change-driven shifts in environmental conditions alter host distributions and expand suitable habitats for ticks, allowing them to spread into previously unaffected regions^5,6^. The expansion of tick populations and the associated risks of pathogen transmission across species make a One Health approach essential^3^. Efforts to document tick distribution have largely relied on tick occurrence records, without integrating related biological, ecological, and tick management data into accessible platforms. For instance, Zhang *et al*.^7^ compiled a georeferenced dataset of tick species distributions across China, which was subsequently expanded and used to map tick-borne pathogen distributions using machine learning^2^. Similar efforts exist in Europe and North America, through platforms such as the VectorNet Data Portal (https://www.vectornetdata.org/), the European Centre for Disease Prevention and Control (ECDC) Tick Maps (https://www.ecdc.europa.eu/en/disease-vectors/surveillance-and-disease-data/tick-maps), and TickEncounter (https://web.uri.edu/tickencounter/fieldguide/), among others. Comparable efforts remain limited in India, where tick-borne diseases such as Kyasanur Forest Disease and Crimean-Congo Hemorrhagic Fever present significant public health risks^8^. Additionally, tick infestations impose a considerable economic burden on the livestock industry through reduced productivity and disease transmission, particularly in cattle and buffalo^9^. Furthermore, India’s considerable geographic diversity, the underreporting of tick-borne diseases in the country, and growing acaricide resistance present substantial challenges for comprehensive surveillance and distribution mapping^9,10^.

Nonetheless, information on tick occurrence, distribution, host associations, and management remains dispersed across the published literature^10,11^. This fragmented nature of data limits its utility for distribution mapping, disease risk assessment, and surveillance efforts. To address this gap, this study presents TickMapKB (https://cb.imsc.res.in/tickmapkb/), a spatial knowledgebase that integrates curated information specific to India on 72 tick species, 647 georeferenced occurrence records, 53 associated pathogens, 122 acaricides with usage information, and 3443 tick-associated proteins, along with morphological keys, within a single, openly accessible, and interoperable platform.

## Methods

### Compilation and curation of data on tick species in India

The overall workflow for constructing TickMapKB, including literature curation and subsequent integration of annotations from other resources, is shown in **Figure 1**. As a first step, relevant studies on tick species and their distribution across India were systematically identified using PubMed. PubMed (https://pubmed.ncbi.nlm.nih.gov/) was used as the primary search database for its open programmatic access without institutional or commercial restrictions, which was necessary for building a reproducible pipeline. The literature search was last conducted on 1 December 2025 using the following query:

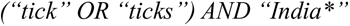

**Figure 1.**
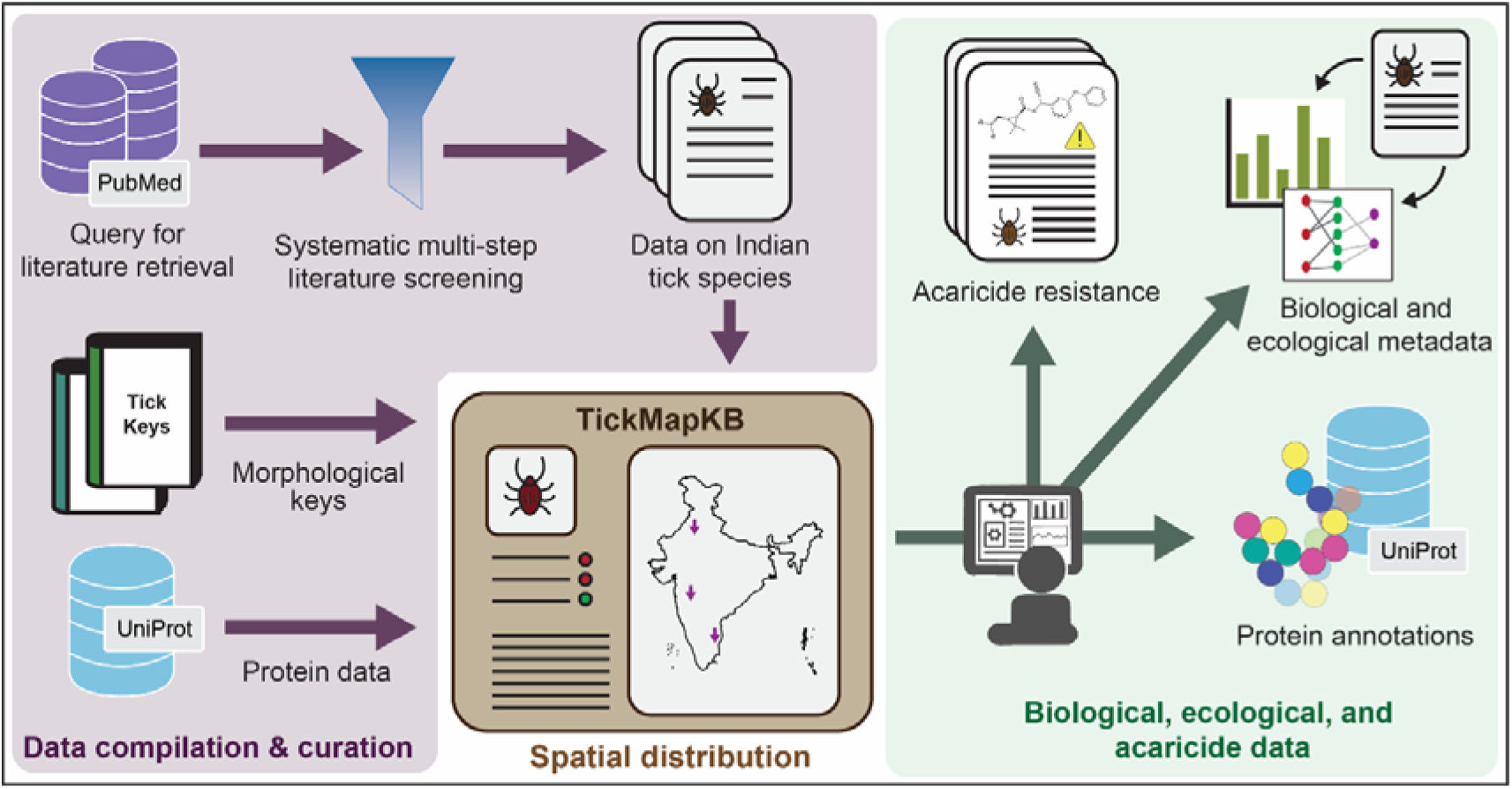
Schematic workflow for the compilation and curation of tick species data in India, integration of annotations from external resources, and incorporation into the TickMapKB web interface.

This literature search retrieved 1464 unique research articles. During the screening stage, articles were retained only if they reported tick occurrence data from India. Additionally, articles published in languages other than English, review articles, laboratory-based studies, and tick-rearing studies were excluded. This process resulted in the removal of 810 articles. The full texts of the remaining 654 articles were then assessed for data curation, of which 264 contained information on tick species and their sampling locations in India and were incorporated into the final dataset. The process for curation of relevant literature is summarized in a PRISMA statement^12^ (**Figure 2**).

**Figure 2.**
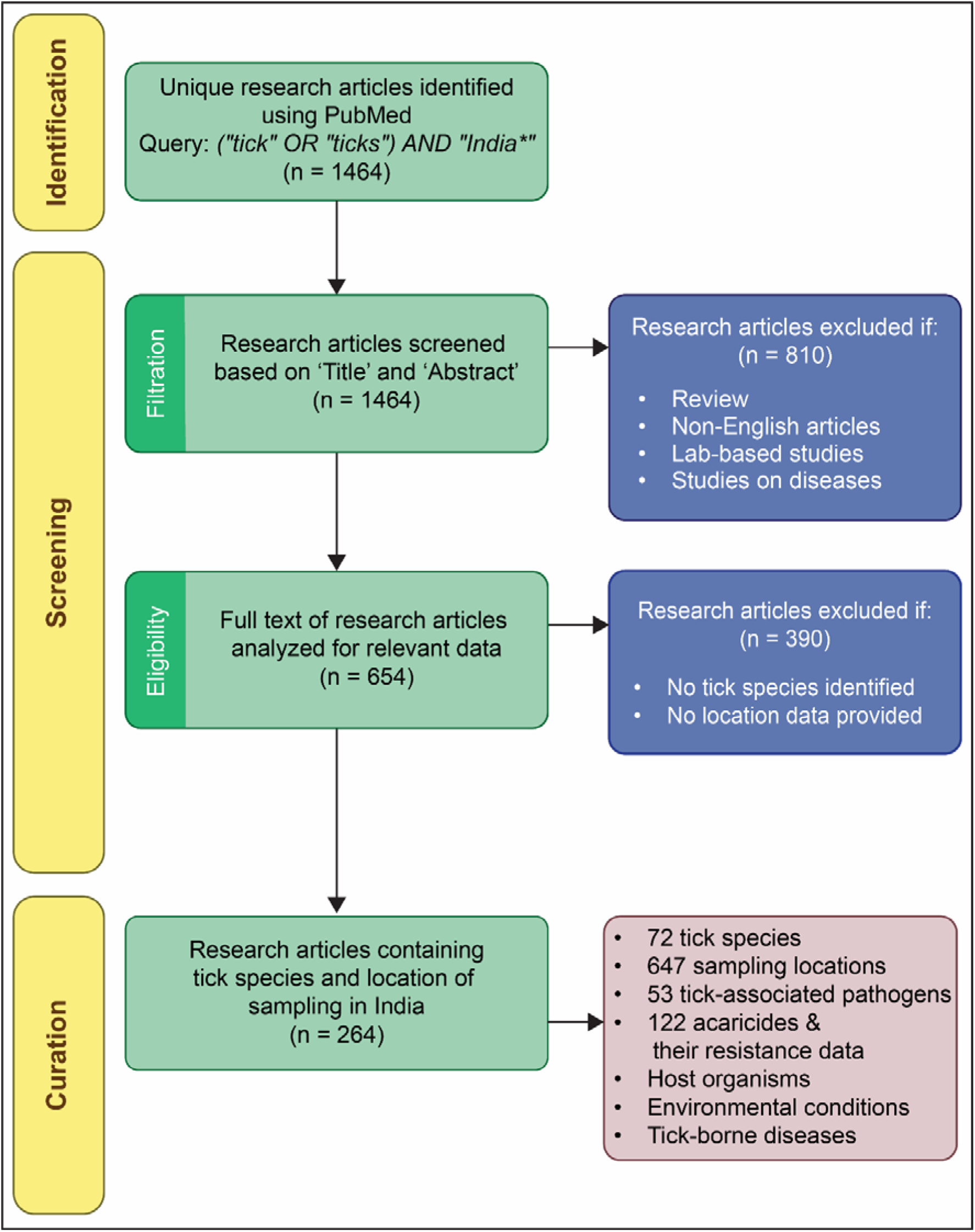
A PRISMA statement summarizing the literature screening and selection process used to identify publications reporting tick occurrence data in India.

Next, from these 264 articles, data were manually extracted on tick species and their geographic sampling locations, including area, district, and state, latitude and longitude coordinates, if provided. Additional biological data were extracted, including host organisms, tick collection methods, tick identification methods, tick-associated pathogens, tick-associated diseases, acaricides, non-chemical tick control methods, and the environmental conditions under which ticks were sampled. Further, acaricide resistance data were recorded as LC50 values, resistance factors, and resistance levels^13^, along with the geographic location of the sampling site. All data were curated by one team member and reviewed by a second, with final verification being done by domain experts.

This effort resulted in the identification of 72 tick species belonging to 10 genera, of which 7 were classified only at the genus level. The genus *Haemaphysalis* had the highest number of species (30), followed by *Rhipicephalus* (12) and *Hyalomma* (11) (**Figure S1A**), but *Rhipicephalus* was associated with the majority of the occurrence records (**Figure S1B**). These species corresponded to 647 unique geographical occurrence sites (**Figure 3**). Moreover, 53 pathogens associated with the curated tick species were identified. The pathogens Kyasanur Forest Disease virus, *Rickettsia spp.*, and *Coxiella burnetii* were associated with the highest number of tick species (**Figure S2A**), while goats and cattle were the most frequently reported host organisms across tick species (**Figure S2B**). The majority of reported resistance instances occurred at level I, followed by susceptible (S) and level II (**Figure 4**), and were associated with 10 acaricides and four tick species, with the most frequent occurrences recorded for Deltamethrin.

**Figure 3.**
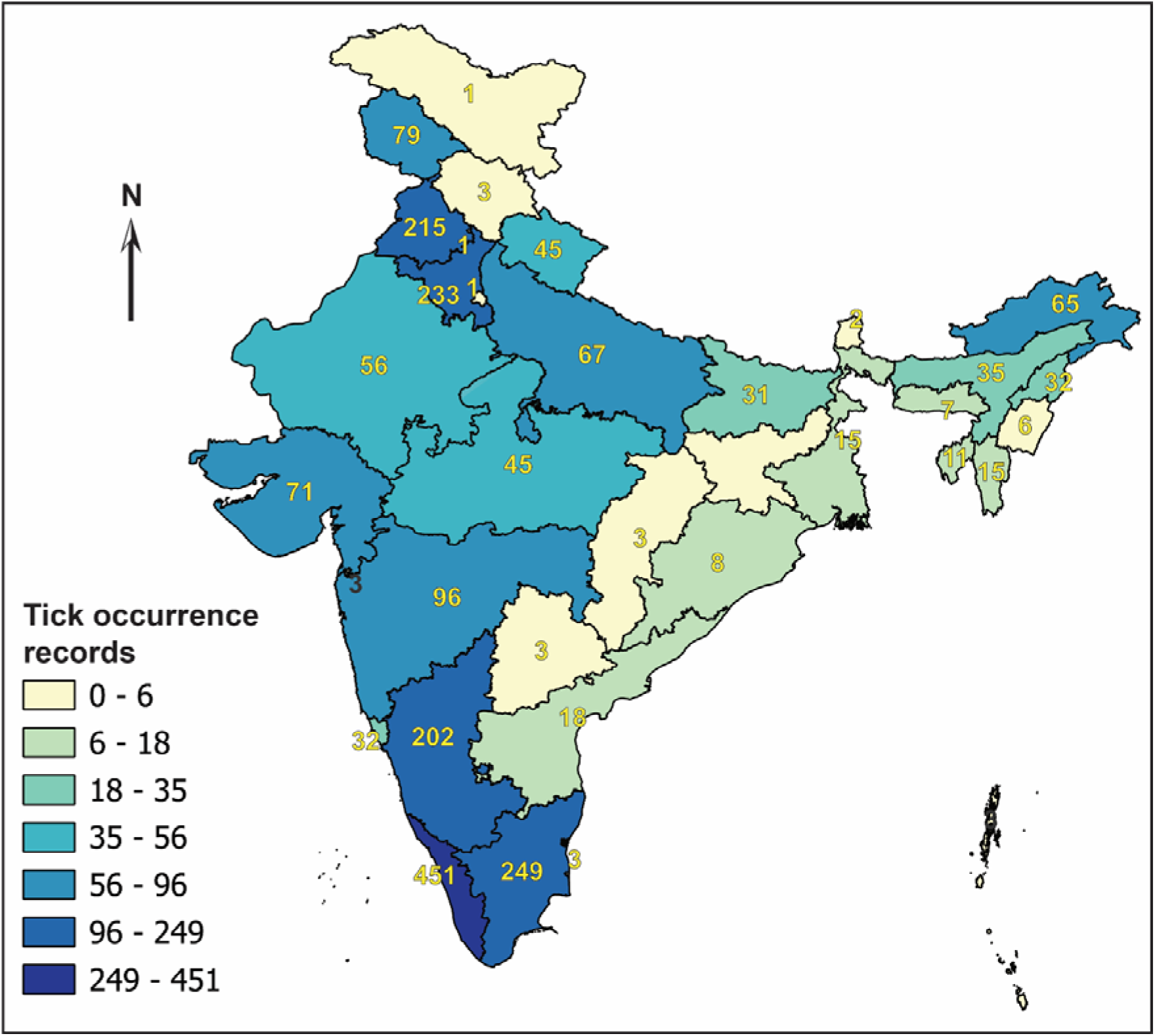
Distribution of tick occurrence records in TickMapKB for India. State shading and labels indicate the total number of occurrence records compiled for each state in TickMapKB.

**Figure 4.**
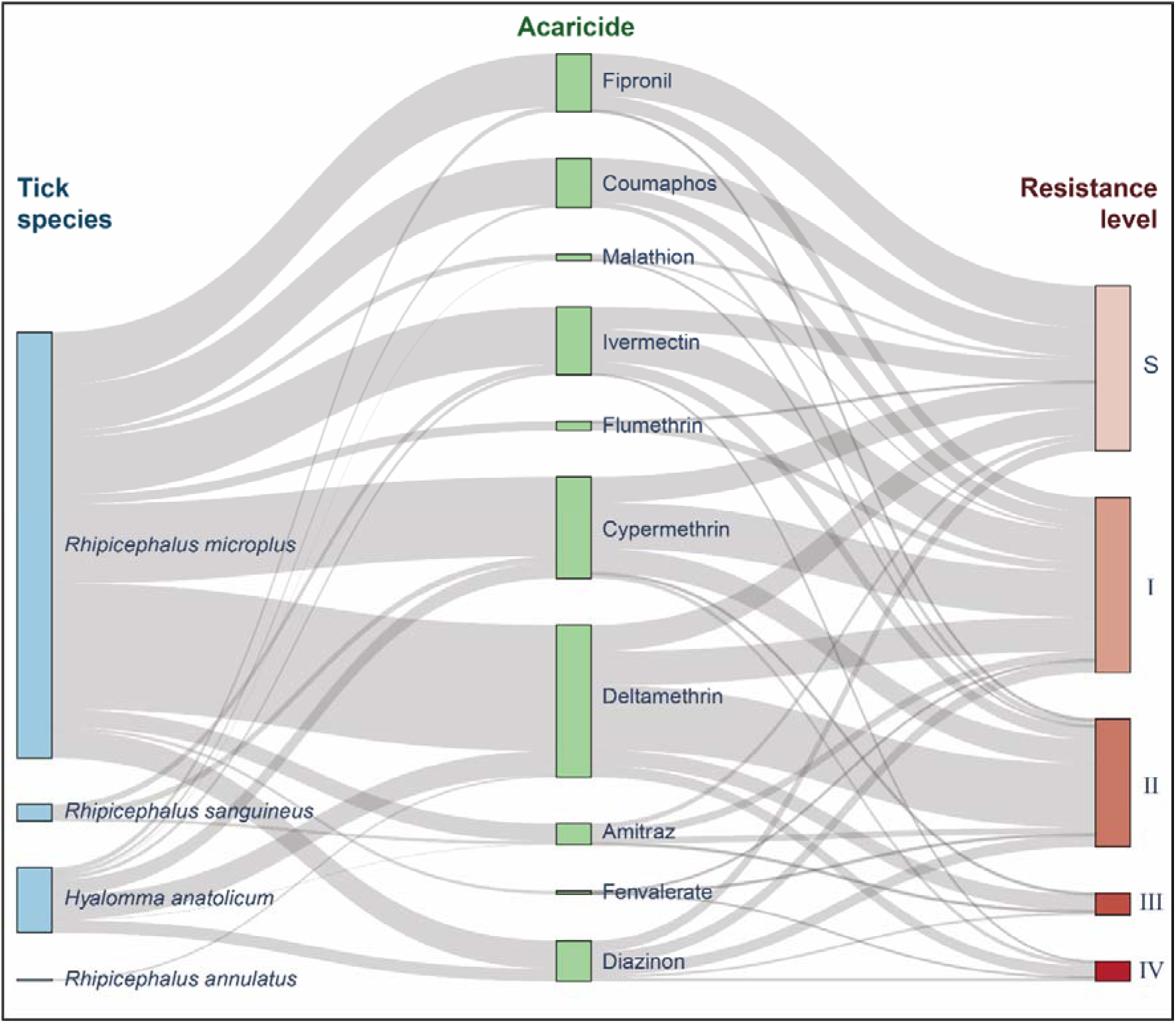
Sankey diagram illustrating the reported acaricide resistance levels in ticks. Flows connect tick species to acaricides and corresponding resistance levels, with the width of each flow representing the number of records.

### Spatial data processing and visualization

Spatial occurrence records were geocoded and visualized using QGIS 3.44.6 (https://qgis.org/), an open-source geographic information system. First, duplicate record entries that share identical values for the reference identifier, tick species, area, district, state, latitude, and longitude were removed. Next, coordinates were standardized by converting those reported in degrees, minutes, and seconds to decimal degrees. The resulting dataset comprised 2107 discrete records corresponding to 647 unique sampling locations.

For records where geographic coordinates were not explicitly reported in the source article, latitudes and longitudes were resolved through geocoding. Geocoding was performed using the MMQGIS plugin (https://plugins.qgis.org/plugins/mmqgis/) in QGIS. The plugin queries the Nominatim geocoding service (https://nominatim.org/) built on OpenStreetMap data to convert place names into geographic coordinates.

To generate interactive distribution maps for tick species recorded in India, vector shapefiles were obtained from the Survey of India (https://surveyofindia.gov.in/) as the base administrative boundary layer, ensuring that national and administrative boundaries were derived from official Indian government sources. First, within QGIS, state and district boundary layers were loaded along with the georeferenced occurrence data. Next, to prevent overlapping records with identical coordinates, point clustering was applied. Finally, the maps were exported as interactive HTML files using the QGIS2Web plugin (https://github.com/qgis2web/qgis2web), with spatial clustering enabled within the layer configurations. These maps are publicly accessible on the TickMapKB web server (https://cb.imsc.res.in/tickmapkb/).

### Data standardization and annotation

The tick species names were standardized by manually mapping them to the corresponding NCBI taxonomy identifiers (https://www.ncbi.nlm.nih.gov/datasets/taxonomy/browser/) and, where available, to the Global Biodiversity Information Facility (GBIF; https://www.gbif.org/) taxonomy identifiers and Barcode of Life Data (BOLD; https://boldsystems.org/) taxonomy identifiers. Of the 72 tick species included in the database, 49 were mapped to both NCBI and BOLD taxonomy identifiers, while 60 were mapped to GBIF identifiers. Subsequently, the curated acaricides were mapped to corresponding PubChem compound identifiers (https://pubchem.ncbi.nlm.nih.gov/) and Chemical Abstracts Service Registry Numbers obtained from CAS Common Chemistry (https://commonchemistry.cas.org/). Structural information was retrieved for 40 of the 122 acaricides, whereas the remaining entries consisted primarily of plant extracts and other natural products for which standardized chemical structures were not available.

Additionally, to facilitate tick identification, morphological keys, which are observable physical characters to differentiate and classify organisms to the species level, were compiled from 24 published sources. Keys were collected for adult males and females, as well as for nymphal and larval stages where available. Further, protein annotations were retrieved by querying species-specific NCBI taxonomy identifiers against the UniProt database^14^ (https://www.uniprot.org/) using the UniProt REST API^15^ (https://www.uniprot.org/api-documentation/uniprotkb) and the Requests library (version 2.32) in Python. This process yielded 3443 protein records across 37 tick species, together with associated UniProt accession identifiers, functional annotations, Protein Data Bank (PDB) structures, subcellular localization information, and Gene Ontology (GO) biological process, cellular component, and molecular function annotations (**Figure 5**).

**Figure 5.**
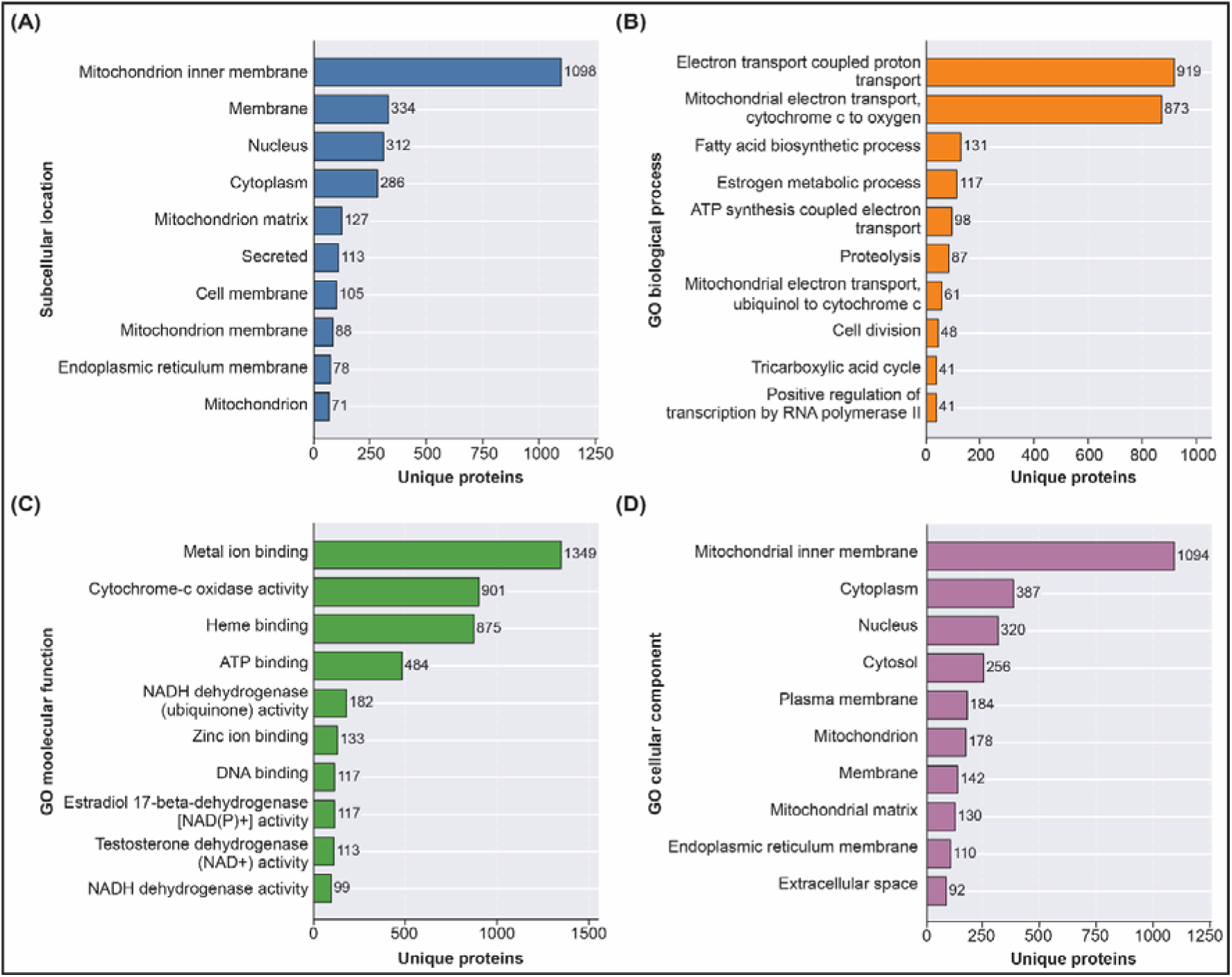
Distribution of protein annotations in TickMapKB. Bar plots show the ten most frequently represented: (A) subcellular localization terms, (B) Gene Ontology (GO) biological process, (C) GO molecular function, and (D) GO cellular component annotations among unique tick proteins.

### Implementation of the TickMapKB web interface and database

The curated datasets in this study were integrated into TickMapKB (https://cb.imsc.res.in/tickmapkb/), a publicly accessible spatial knowledgebase for exploring tick distribution, proteomic annotations, acaricides, and acaricide resistance within India. The resource integrates species-level occurrence records, interactive maps, curated morphological identification keys, protein annotations derived from UniProt, and information on acaricides and resistance profiles onto a single platform. The resource includes a dedicated help section with use cases and video tutorials to support data exploration and retrieval. All records are accessible through a searchable web interface and can be downloaded in standard formats without registration. To support the FAIR (Findable, Accessible, Interoperable, and Reusable) data principles^16^, species and chemical records are cross-linked to established external repositories and assigned persistent identifiers where available. All database content, except for images obtained from third-party sources, is distributed under the Creative Commons Attribution 4.0 International (CC BY 4.0) license (https://creativecommons.org/licenses/by/4.0/).

All curated records are maintained in a MariaDB (https://mariadb.org/) database and accessed via Structured Query Language (SQL) queries. The web interface was implemented using PHP (http://php.net/), HTML, CSS, Bootstrap 5 (https://getbootstrap.com/docs/5.0/), and jQuery (https://jquery.com/), and was deployed on an Apache HTTP server (https://httpd.apache.org/) running on a Debian 9.4 operating system. Interactive maps were generated in QGIS and exported using the QGIS2Web plugin, which utilizes the Leaflet JavaScript library (https://leafletjs.com/) for web-based visualization. Additional data visualizations throughout the platform were implemented using the Chart.js JavaScript library (https://www.chartjs.org/).

## Data records

The curated dataset is interactively accessible via the TickMapKB web interface at https://cb.imsc.res.in/tickmapkb/. Moreover, the curated dataset is also available as an Excel workbook comprising four datasheets for which a detailed description is provided below.

### Tick species distribution and biological data

The primary datasheet (**Table S1**) contains 2299 records of curated tick species occurrence and associated biological data across India. Each record corresponds to an occurrence of a tick species at a geographic sampling location (i.e., area, district, state, latitude, and longitude) as reported in the source literature. Further, it includes biological metadata such as host organisms, tick collection and identification methods, associated pathogens and diseases, acaricide and non-chemical tick control methods, and environmental conditions at the time of sampling.

### Morphological keys

The second datasheet (**Table S2**) provides morphological keys for tick identification. The dataset links each tick species in the database to morphological keys curated from 24 literature sources, organized by life stage and sex.

### Acaricide resistance

The third datasheet (**Table S3**) contains 984 records of acaricide resistance, which were extracted from the published literature on tick species reported to be resistant to acaricides in India. For each record, resistance is quantified using LC50 values, along with resistance factors and levels, as reported in the source literature, with resistance levels following the classification criteria of Kumar *et al*. (https://doi.org/10.1016/j.vetpar.2011.04.030) where applicable. Each record is further linked to its geographic sampling location (i.e., area, district, state, latitude, and longitude) and annotated with chemical identifiers, including PubChem Compound Identifiers and Chemical Abstracts Service Registry Numbers.

### Tick-associated protein annotations

Finally, the fourth datasheet (**Table S4**) contains 3443 curated protein records for tick species recorded in the database, sourced from UniProt. For each record, the table provides protein and gene identifiers, reviewed status, subcellular localization, GO annotations across biological process, molecular function, and cellular component, and PDB identifiers, where available.

## Technical validation

The taxonomic and spatial data were validated through multiple manual checks. First, species names in older publications were updated to their current accepted synonyms, where applicable. For instance, records of *Haemaphysalis centropi* were updated to *Haemaphysalis doenitzi*, as Indian specimens originally identified as *H. centropi* by Kohls in 1949 (https://doi.org/10.2307/3273429) were subsequently synonymized with *H. doenitzi* on the basis of shared morphological characteristics (https://doi.org/10.1016/B978-0-12-387811-3.00001-2). Furthermore, reports of *Amblyomma americanum* (lone star tick) and *Ixodes scapularis* (deer tick) in India were considered possible misidentifications, and the corresponding records were excluded. *A. americanum* is native to North America, with its core distribution concentrated in the southeastern states of the United States of America, where it is considered a medically significant tick species (https://doi.org/10.1371/journal.pone.0209082; https://doi.org/10.1146/annurev.ento.48.091801.112728). Similarly, *I. scapularis* is also native to North America, ranging from the southern part of Canada through the eastern part of the United States of America, where it is a known vector of Lyme disease (https://doi.org/10.1186/s13071-015-1185-7; https://doi.org/10.1016/j.lana.2024.100706). Finally, records lacking species-level identification were reviewed by a taxonomist, who assigned species names based on morphological features visible in images within the respective articles, wherever possible.

Further, older names of places (geographic locations) were updated to reflect recent administrative changes, such as Allahabad, which was renamed Prayagraj. Also, typographical errors in location names were corrected. Moreover, it should be noted that the spatial resolution varied across published articles. Hence, when a study provided only a broader district or state name without an exact sampling location or its coordinates, the location was resolved to the corresponding administrative capital. Moreover, the coordinates obtained from MMQGIS-based geocoding were manually verified against Google Earth (https://earth.google.com/web; accessed in April 2026) to ensure spatial accuracy.

## Usage notes

TickMapKB integrates curated information on tick occurrence within India, along with associated pathogens, morphological keys, acaricide resistance profiles, and protein data into a single and interactive spatial knowledgebase. A comparison of existing tick resources across data types and functionalities (**Figure 6**) revealed that most are region-specific, covering Europe, North America, or China, with none specific to India. These include curated resources such as a dataset by Zhang *et al*.^7^ for China, which includes occurrence records for 123 species, VectorNet (https://www.vectornetdata.org/) and ECDC Tick Maps (https://www.ecdc.europa.eu/en/disease-vectors/surveillance-and-disease-data/tick-maps) for Europe, and TickEncounter (https://web.uri.edu/tickencounter/fieldguide/) and VectorSurv (https://maps.vectorsurv.org/arbo) for North America. Apart from literature-curated and surveillance-based platforms, citizen science initiatives such as Tekenradar (https://www.tekenradar.nl/), with over 60000 tick bite reports, and eTick (https://www.etick.ca/etickapp/en/ticks/public/map), which provides expert identification within two days, demonstrate the value of continuous community-driven updates. Other resources, such as the University of Texas Medical Branch (UTMB) Tick Map (https://www.utmb.edu/wgcvbd/tick-map) and Zecke Tique Tick (https://zecke-tique-tick.ch/en/tickbite-map-switzerland/), are similarly constrained to specific regions, covering Texas and Switzerland, respectively. While most of these resources are limited to occurrence data and interactive maps, the UTMB Tick Map and TickEncounter additionally provide species information, and TickEncounter and VectorSurv also include pathogen- and disease-related data. However, none of these resources cover India. TickMapKB thus provides an India-specific platform spanning a range of data types and functionalities not covered by other existing resources (**Figure 6**).

**Figure 6.**
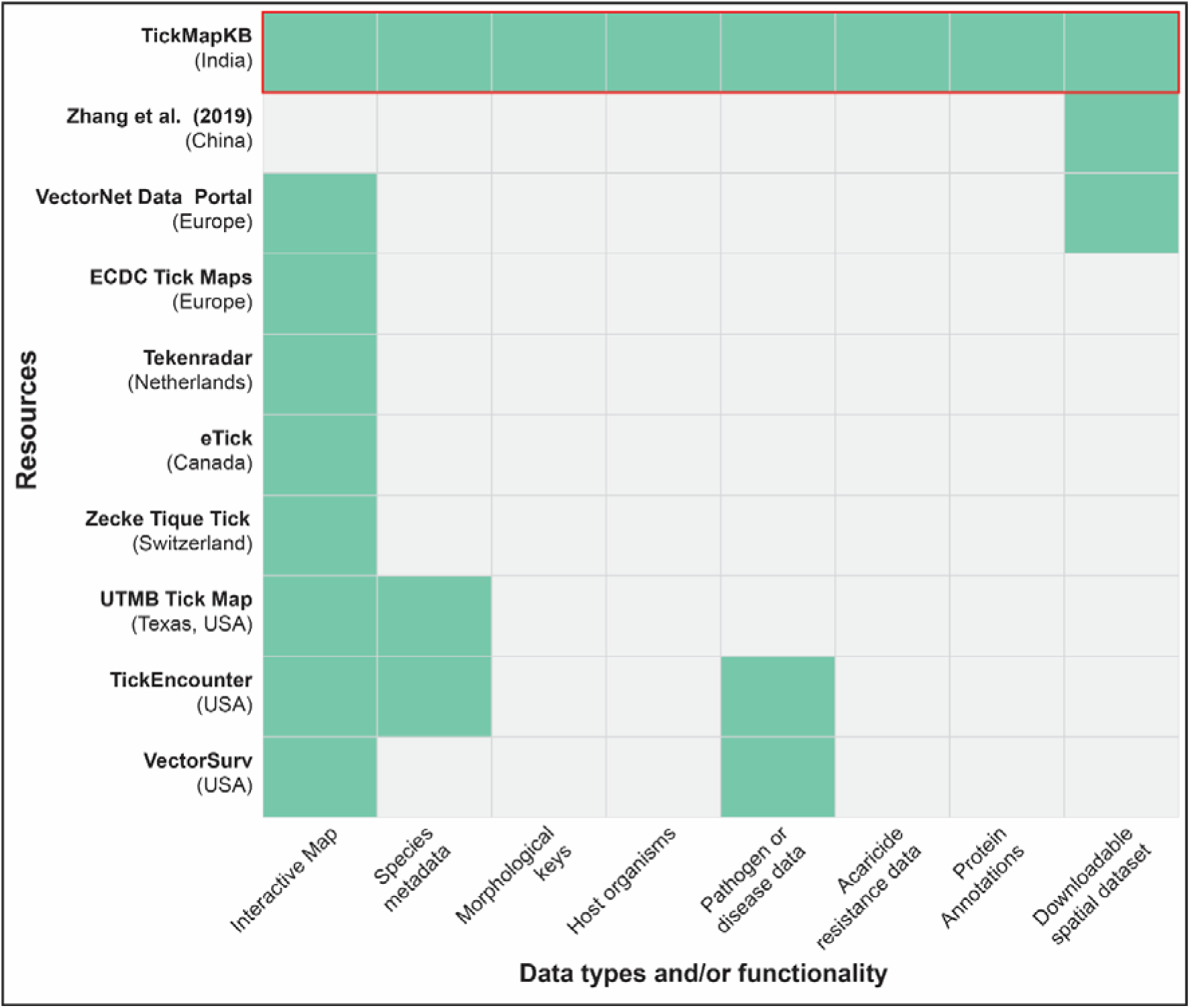
Feature comparison of TickMapKB (highlighted with a red border) with existing tick-related resources. Each row represents a resource with geographic coverage indicated in parentheses, and each column denotes available data type and/or functionality. Filled cells indicate the presence of a feature.

It is worthwhile to note that the species and occurrence count in TickMapKB reflect what is currently documented in the published literature and are likely an underestimate, given known underreporting and uneven geographic sampling across India. Further, the acaricide resistance data across studies are not directly comparable, as no established susceptible reference isolate exists for Indian tick populations. Therefore, individual studies differ in their calculation of resistance factors and levels. Similarly, tick identification methods varied across published studies, which may affect the reliability of some species-level records.

Nonetheless, the resource will support a broad range of users across diverse applications such as tick identification, disease surveillance, vector control, and potentially drug or vaccine target discovery. A detailed description of these use cases is available on the Help page of the TickMapKB web interface (http://cb.imsc.res.in/tickmapkb/help#use-cases), and selected examples are discussed below.

### Tick species identification

Researchers working with tick specimens can use TickMapKB for species identification. If the genus is already known, species pages can be browsed directly to compare morphological key entries across life stages and sexes. Where the genus is unknown but morphological characteristics are observable, the ‘Advanced Search’ interface allows key-based queries across the dataset, returning candidate species matches that can then be cross-checked against taxonomy and known host data.

### Tick-borne disease surveillance

TickMapKB supports region-specific querying of tick species and their associated pathogens. Users can filter occurrence records by district or state to identify tick species documented in a given area, and navigate to species-level pathogen and disease data to potentially prioritize diagnostic testing strategies.

### Acaricide Selection and Resistance Assessment

Users can potentially use the acaricide resistance profiles to evaluate the efficacy of available compounds against specific tick species in a given region. Resistance data, including LC50 values, resistance factors, and resistance levels, can be filtered by species and state to examine patterns of acaricide resistance.

### Tick protein data for drug target discovery

The protein data can aid in the identification of potential drug and vaccine targets through proteome-level exploration of tick species. For instance, secreted proteins exposed to the host immune system during feeding can be filtered and further refined using keyword annotations to identify candidate targets of interest. Interoperability with UniProt and the PDB allows direct follow-up for sequence, functional, and structural information. Furthermore, genus-level filtering also allows comparison of protein profiles across related species, which may reveal conserved protein families with shared biological roles, such as host immune evasion.

## Data availability

The curated data associated with this study are included in the article or are publicly accessible via the associated website: https://cb.imsc.res.in/tickmapkb/.

## Author contributions

Conceptualization: Shreyes Rajan Madgaonkar, Srikanth Srirama, Areejit Samal; Methodology: Shreyes Rajan Madgaonkar, Shrish Vashishth, Elango Ayyanar, Srikanth Srirama, Areejit Samal; Data curation: Shreyes Rajan Madgaonkar, Shrish Vashishth; Formal analysis: Shreyes Rajan Madgaonkar, Shrish Vashishth, Elango Ayyanar, Srikanth Srirama, Areejit Samal; Visualization: Shreyes Rajan Madgaonkar, Shrish Vashishth, Software: Shreyes Rajan Madgaonkar, Shrish Vashishth; Writing – original draft: Shreyes Rajan Madgaonkar, Shrish Vashishth, Elango Ayyanar, Srikanth Srirama, Areejit Samal; Writing – review & editing: Shreyes Rajan Madgaonkar, Shrish Vashishth, Elango Ayyanar, Srikanth Srirama, Areejit Samal; Supervision: Areejit Samal

## Supplementary Figure Captions

**Figure S1.** Genus-level representation of ticks in TickMapKB. **(A)** Number of unique tick species represented within each genus. **(B)** Number of occurrence records associated with each genus.

**Figure S2.** Heatmaps showing **(A)** the distribution of species among the ten most frequently reported tick-associated pathogens across tick genera, and **(B)** the distribution of species among the ten most frequently reported host organisms across tick genera.

## Supplementary Table Captions

**Table S1**: This table contains the curated data on tick species distribution and associated biological data across India. For each record, the table provides the tick species, family, geographic location (area, district, and state), latitude and longitude coordinates in decimal degrees, host organism(s), tick collection site or method, study period, tick identification method, general environmental condition(s), seasonal condition(s), acaricide, effective acaricide dose, non-chemical prevention method(s), associated pathogens, disease caused, disease-associated adverse effects / symptoms, disease treatment, and the source reference. Acaricide entries are further annotated with PubChem Compound Identifiers (PubChemID) and Chemical Abstracts Service Registry Numbers (CASRN). The following fields may contain multiple pipe-separated (‘|’) entries: host organism(s), tick identification method, general environmental condition(s), seasonal condition(s), non-chemical prevention method(s), associated pathogens, disease caused, disease-associated adverse effects / symptoms, and disease treatment.

**Table S2**: This table contains the morphological identification keys compiled for tick species recorded in the database. For each entry, the table provides the tick species, life stage and sex (male, female, nymph, or larva), the morphological key used for identification, and the source reference.

**Table S3**: This table contains the curated acaricide resistance data for tick species recorded in the database. For each record, the table provides the tick species, life stage, geographic location (area, district, and state), latitude and longitude coordinates in decimal degrees, acaricide, test type, LC50 value and its unit, resistance factor / ratio, resistance level, and the source reference. Acaricide entries are further annotated with PubChem Compound Identifiers (PubChemID) and Chemical Abstracts Service Registry Numbers (CASRN). Resistance levels are reported as recorded in the source literature, where applicable following the classification criteria described in Kumar *et. al.* (2011) (https://doi.org/10.1016/j.vetpar.2011.04.030): susceptible (RF ≤ 1.4), level I (RF = 1.5-5.0), level II (RF = 5.1-25.0), level III (RF = 26-40), and level IV.

**Table S4**: This table contains the protein data for tick species in the database curated from UniProt. For each entry, the table provides the tick species, NCBI Taxonomy Identifier (TaxID), UniProt accession number, protein name, gene name, protein length in amino acids (aa), reviewed status, subcellular location, Gene Ontology (GO) annotations for biological process, molecular function, and cellular component, associated keywords, Protein Data Bank identifier, and a brief functional description of the protein.

