## Supplementary material for "TickMapKB: A FAIR Spatial Knowledgebase of Tick Species and Associated Pathogens in India": Figure S

### Supplementary Figures S1-S2

**for**

### TickMapKB: A FAIR Spatial Knowledgebase of Tick Species and Associated Pathogens in India

Shreyes Rajan Madgaonkar[a,b], Shrish Vashishth[a], Elango Ayyanar[c], Srikanth Srirama[c], Areejit Samal[a,b,*]

[a]*The Institute of Mathematical Sciences (IMSc), Chennai, India*

[b]*Homi Bhabha National Institute (HBNI), Mumbai, India*

[c]*ICMR-National Institute for Vector Control Research, Puducherry, India*

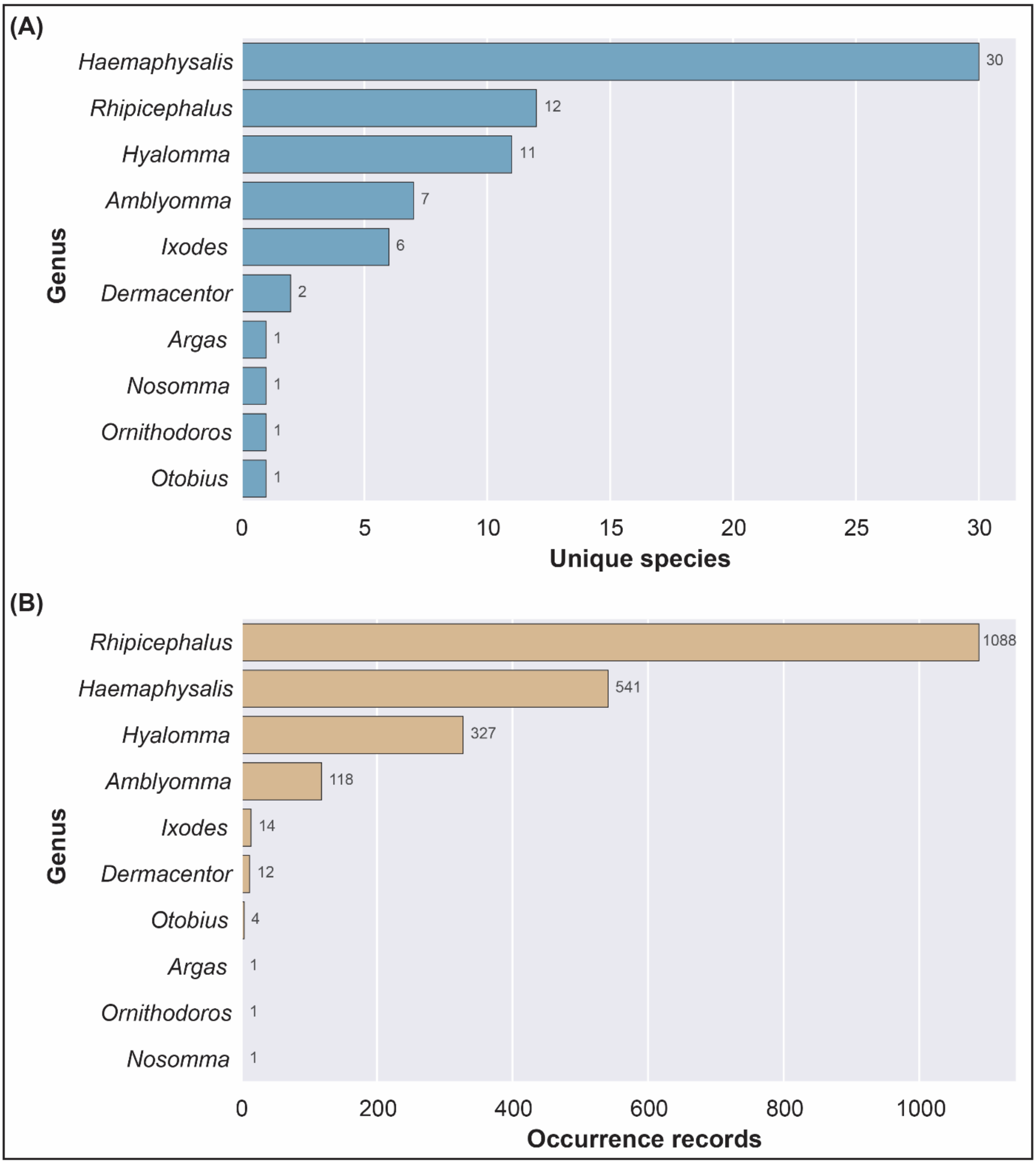


**Figure S1**. Genus-level representation of ticks in TickMapKB. **(A)** Number of unique tick species represented within each genus. **(B)** Number of occurrence records associated with each genus.

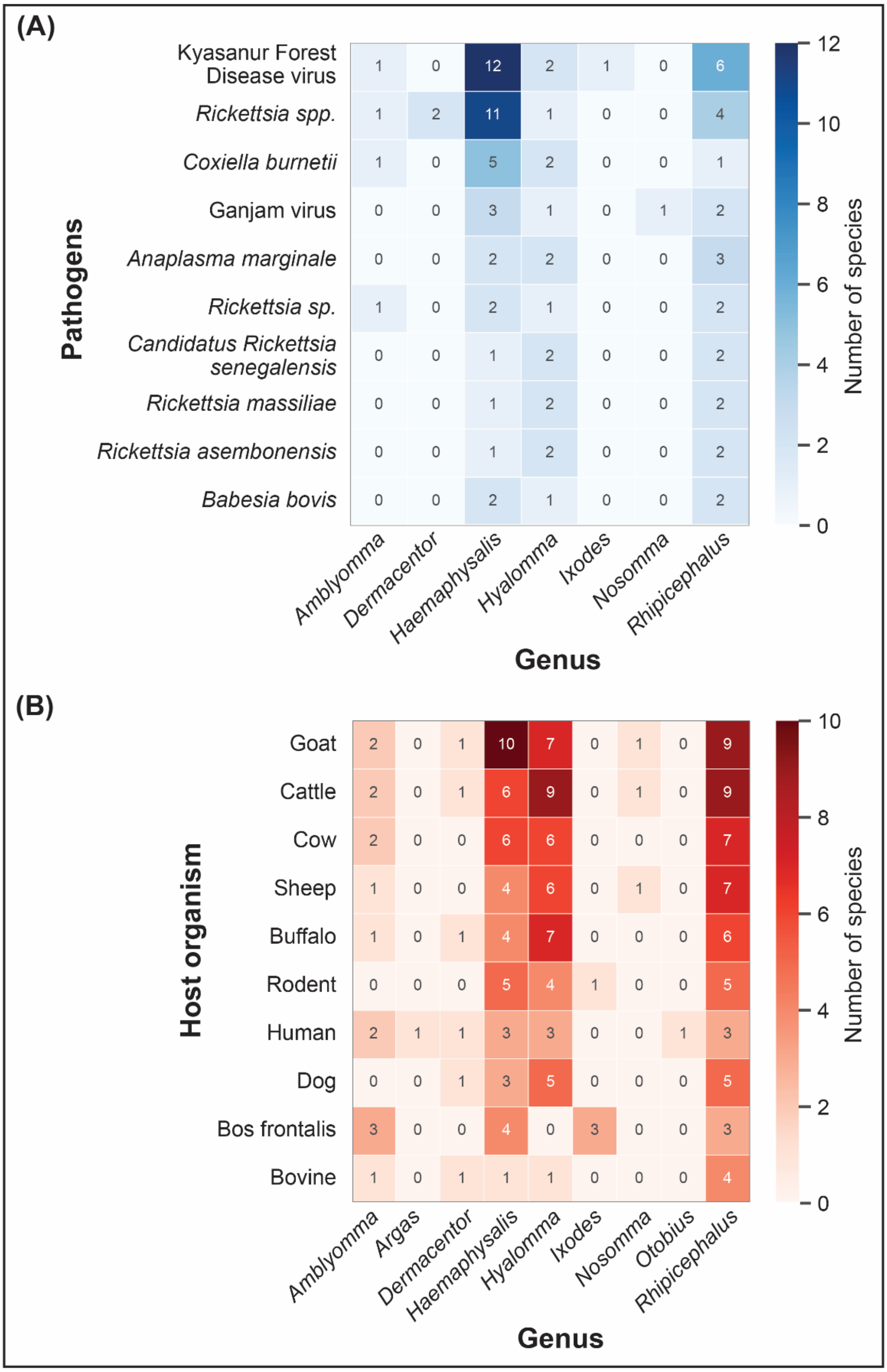


**Figure S2**. Heatmaps showing **(A)** the distribution of species among the ten most frequently reported tick-associated pathogens across tick genera, and **(B)** the distribution of species among the ten most frequently reported host organisms across tick genera.
